# DMBT1 promotes SARS-CoV-2 infection and its SRCR-derived peptide inhibits SARS-CoV-2 infection

**DOI:** 10.1101/2025.03.06.641801

**Authors:** Chenxi Zhu, Ziqiao Wang, Zhendong Pan, Xinjia Mai, Yiyun Chen, Wen Zhang, Ping Zhao, Hailin Tang, Rong Zhang, Dapeng Zhou

## Abstract

DMBT1 is a large scavenger receptor cysteine rich (SRCR) B protein that has been reported as a tumor suppressor gene and a co-receptor for HIV-1 infection. Here we found DMBT1 is a major mucosal protein bound to SARS-CoV-2. Overexpression of DMBT1 in 293T cells enhanced infection by SARS-CoV-2 in ACE2 dependent manner. Blocking experiments using overlapping peptide library of SRCR domain of DMBT1 showed that CQGRVEVLYRGSWGTV peptide, which contains bacteria-binding VEVLXXXXW motif, could inhibit SARS-CoV-2 infection. High concentration of the peptide can significantly inhibit the replication of SARS-CoV-2 in hamsters. Single cell sequencing analysis showed the highest expression abundance of DMBT1 at airway submucosal glands. These results demonstrate that membrane DMBT1 can promote SARS-CoV-2 infection, and serve as a candidate target for antiviral development.

**Author Summary:** We discovered that a protein named as DMBT1, which is found in our mucous membranes, plays a significant role in SARS-CoV-2 infection. DMBT1 is a large glycoprotein that has previously reported linked to tumor suppression and HIV-1 infection. Our single-cell sequencing analysis revealed that DMBT1 is most abundant in the airway submucosal glands. In our study, we found that when DMBT1 is overexpressed in cells, it enhances SARS-CoV-2 infection. Using a peptide library from DMBT1, we pinpointed a specific peptide, CQGRVEVLYRGSWGTV, which can inhibit SARS-CoV-2 infection. At high concentrations, it significantly reduces the virus’s ability to replicate in hamsters. These findings suggest that DMBT1 on the cell membrane can promote SARS-CoV-2 infection and highlight DMBT1 as a potential target for developing antiviral treatments.

## Introduction

Several receptors have been reported for SARS-CoV-2 responsible for COVID-19. SARS-CoV-2 primarily utilizes the ACE2 receptor for host cell entry through TMPRSS2-mediated membrane fusion, or cathepsin L (CTSL)-mediated endosomal membrane fusion[1-3]. Viral attachment and infection involve heparan sulfate dependent binding to ACE2[4]. In addition, LDLRAD3, TMEM30A, and CLEC4G have been reported to effectively mediate viral invasion into cells in an ACE2 independent manner[5]. Loss of CD147 or blocking with antibody would inhibit SARS-CoV-2 amplification, and CD147 could mediate virus entry into host cells through endocytosis^[6]^. ASGR1 and KREMEN1 can interact with multiple Spike protein domains and can directly mediate ACE2 independent SARS-CoV-2 infection, but have no effect on SARS and MERS infection[7]. TMEM106B, a lysosomal transmembrane protein, interacts synergistically with the receptor heparan sulfate to mediate the entry of SARS-CoV-2 into ACE2 negative cell lines[8].

DMBT1, also known as GP340 (DMBT1^GP340^) or salivary agglutinin (DMBT1^SAG^), is a glycoprotein with 8 to 13 continuously repetitive cysteine rich scavenger receptor (SRCR) sequences, with genetic polymorphisms and complex surface glycosylation modifications[9, 10]. DMBT1 binds to a wide range of bacteria and viruses as part of the innate defense of mucosal surfaces. DMBT1 has been implicated in interactions with both HIV-1 and influenza A virus (IAV). DMBT1 in lung and saliva suppresses hemagglutination activity and infectivity of IAV by mediating viral particle aggregation through calcium-independent interactions with sialylated glycans[11, 12] Initially, DMBT1^SAG^ and the recombinant N-terminal SRCR-SID (SRCR interspersed domain) of DMBT1 were shown to bind the HIV-1 envelope glycoprotein gp120, thereby inhibiting HIV-1 infection[13, 14]. However, DMBT1 expressed on genital tract epithelia promotes HIV-1 binding and transcytosis via specific gp120 protein-protein interactions[15, 16]. Macrophage-expressed GP340 binds to the HIV-1 envelope and enhances viral fusion and infection[17].DMBT1 on the mucosal surface can aggregate bacteria. The interaction between *Streptococcus mutans* and DMBT1 in the oral cavity has been studied in the context of cariogenesis. DMBT1 induces bacterial aggregation in saliva, resulting in the clearance of bacteria from the mouth, while DMBT1 in the adsorbed form on the tooth surface induces bacterial adhesion, resulting in bacterial accumulation[18-21]. DMBT1 in saliva binds to *Staphylococcus aureus* and may promote colonization in the oral cavity[22]. Reducing the number of N-terminal SRCR domains from 13 to 8 will result in a 20-45% reduction in bacterial binding, including *Streptococcus mutants (IngBritt)*, *Streptococcus gordonii (HG222)*, *Escherichia coli (F7)*, and *Helicobacter pylori (NCTC 11637)*[20]. DMBT1 in tears suppresses the twitching motility of *Pseudomonas aeruginosa* through its N-glycosylation and interaction with bacterial pili, thereby reducing bacterial virulence[23, 24]. The interactions of DMBT1 with various pathogens make it an important regulator at the host-microbe interface, balancing pathogen clearance and colonization.

A peptide sequence in the SRCR domain of DMBT1 has also been shown to bind to *Streptococcus mutans*[25]. The peptide derived from SRCR sequence (QGRVEVLYRGSWGTVC) was named SRCRP2. In this peptide, residues VEVL and W of SRCRP2 are the key sites to binding. DMBT1 recognizes and binds to the leucine-rich repeat (Lrr) domains present on various bacterial surface proteins, such as Spy0843 from *Streptococcus pyogenes*, LrrG from *Streptococcus agalactiae*, and BspA from *Tannerella forsythia*[26]. Peptides derived from the SRCR domain, containing the VEVLXXXXW motif, have been demonstrated to inhibit the binding of recombinant Spy0843 and DMBT1^GP340^. However, it has not yet been reported whether the peptides derived from SRCR domain affect viral infection.

In human respiratory and intestinal cells, less than 10% of cells express both ACE2 and TMPRSS2[27]. These cells can be divided into three types: Goblet secretory cells in the nasal cavity, type II alveolar epithelial cells and absorptive intestinal epithelial cells. The expression of DMBT1 in AT2 cells was also higher than that of CD147[28]. It is worth noting that DMBT1 and CD147 showed a single-cell expression pattern similar to ACE2 in different AT2 subsets, and the expression of ACE2 and DMBT1 in AT2 cells was significantly positively correlated[28]. Higher DMBT1 and lower ACE2 receptor in children may be the reason why they are less susceptible to SARS-CoV-2 infection compared to adults[29, 30]. In addition, SARS-CoV-2 infection promoted alternative splicing to generate 6 new isoforms of DMBT1 in the iPSC-airway organoids at 24 h post-infection, which may be related to virus clearance or antiviral activity[31]. This implies that DMBT1 may have unique functions during viral infection.

We previously analyzed the affinity-proteomics data of saliva absorbed to plate-bound Spike protein of SARS-CoV-2, and identified major virus-binding proteins as MUC7, DMBT1, neutrophil defensins, and MUC5B. We further examined the infection efficiency of SARS-CoV-2 on 293T cells and 293T ACE2 knockout (293T-ACE2KO) cells overexpressing DMBT1. We also synthesized 16 peptides with overlapping amino acid sequences based on SRCR domain and tested their capacity to inhibit SARS-CoV-2 infection. Among them, peptide 7 (CQGRVEVLYRGSWGTV) emerged as the most potent antiviral candidate, demonstrating significant efficacy in both *in vitro* and *in vivo* models. High-dose treatment with peptide 7 significantly inhibits SARS-CoV-2 infection in hamsters. By synthesizing a single amino acid mutated peptide, it was revealed that the cysteine residue in peptide 7 is essential for both dimer formation and antiviral efficacy.

## Result

### DMBT1 enhanced ACE2-dependent SARS-CoV-2 infection

We cloned the DMBT1 gene containing 14 scavenger receptor cysteine-rich (SRCR) domains into pLV-GFP-IRES-puro vector, and overexpressed it in wild-type 293T or 293T-ACE2KO cells by lipofectamine transfection. After puromycin selection, DMBT1 positive cells were sorted by flow cytometry (Fig 1A) and validated through western blot (Fig 1B).

**Fig 1.**
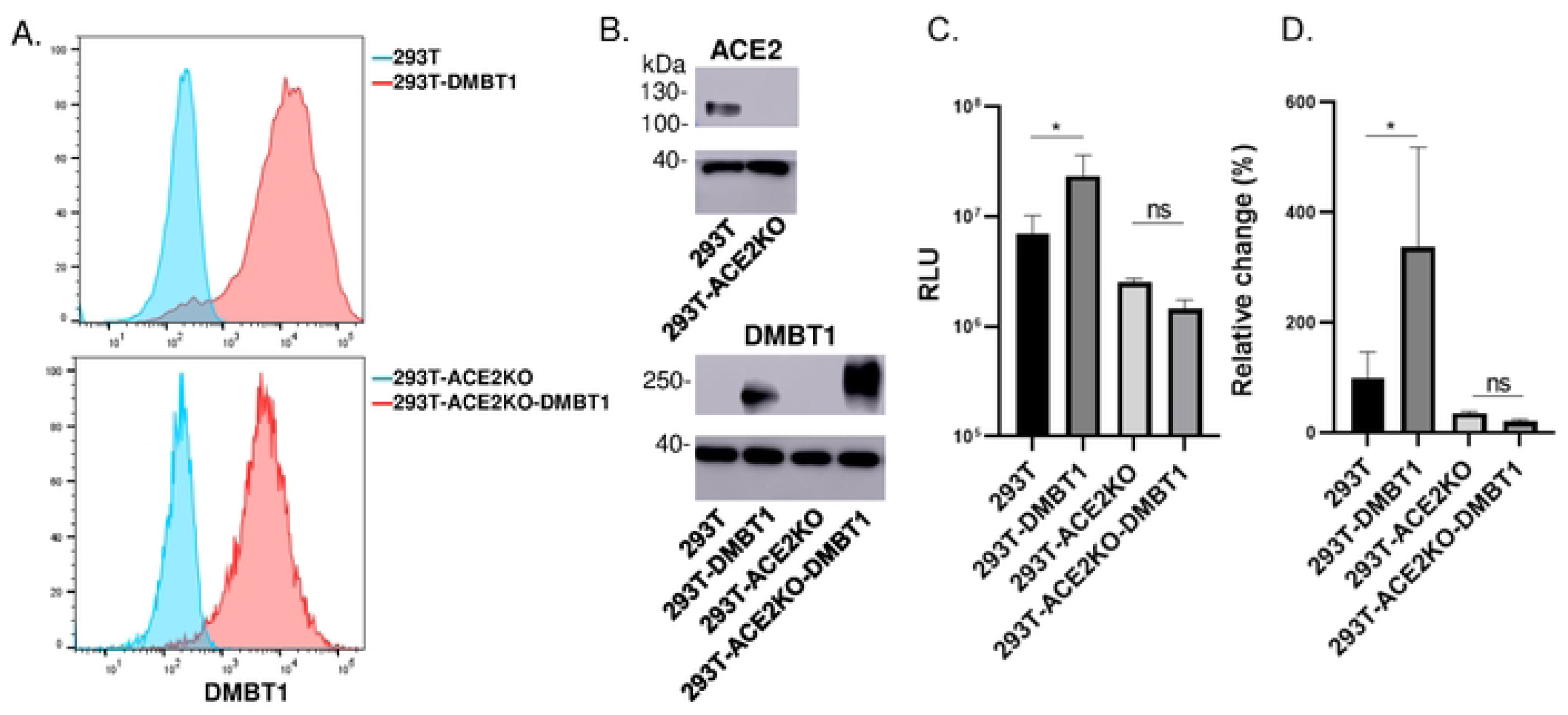
DMBT1 enhanced ACE2 dependent SARS-CoV-2 VLPs infection. (A) Flow cytometry staining of cell lines over-expressing DMBT1. (B) Western blot of 293T and 293T-ACE2KO using anti-ACE2 mAb. Cell lines over-expressing DMBT1 were analyzed using anti-DMBT1 pAb. (C) Luminescence values (RLU) of 293T and 293T-ACE2KO over-expressing DMBT1 cells infected with VLP (MOI=0.03). (D) Relative infection changes of 293T and 293T-ACE2KO cells over-expressing DMBT1. Data were representative of three independent experiments. *, P<0.05; Statistic significance was assessed by one-way ANOVA.

To examine whether DMBT1 enhances viral infection, we first infected the cells with single-cycle transcription and replication-competent SARS-CoV-2 virus-like-particles (trVLP-Nluc), in which the N gene is replaced by NanoLuc luciferase reporter and packaged in N-expressing cells[32, 33]. Overexpression of DMBT1 in 293T-cells showed significant enhancement of infection, which was three times higher than that of 293T wild-type cells (Fig 1C and D). However, overexpression of DMBT1 did not enhance the infection in ACE2-KO 293T cells. These results indicate that DMBT1 can promote virus invasion in the presence of ACE2.

To examine the distribution of DMBT1 expression in human airway cells, we analyzed publicly available scRNA-seq/snRNA-seq data from lung tissues[39] (Fig 2A). Within the airway epithelial cells, the highest expression level and proportion of DMBT1 were observed in submucosal gland (SMG) duct cells and SMG serous cells, with a subset of cells in AT2 and club cells also exhibiting high expression.

**Fig 2.**
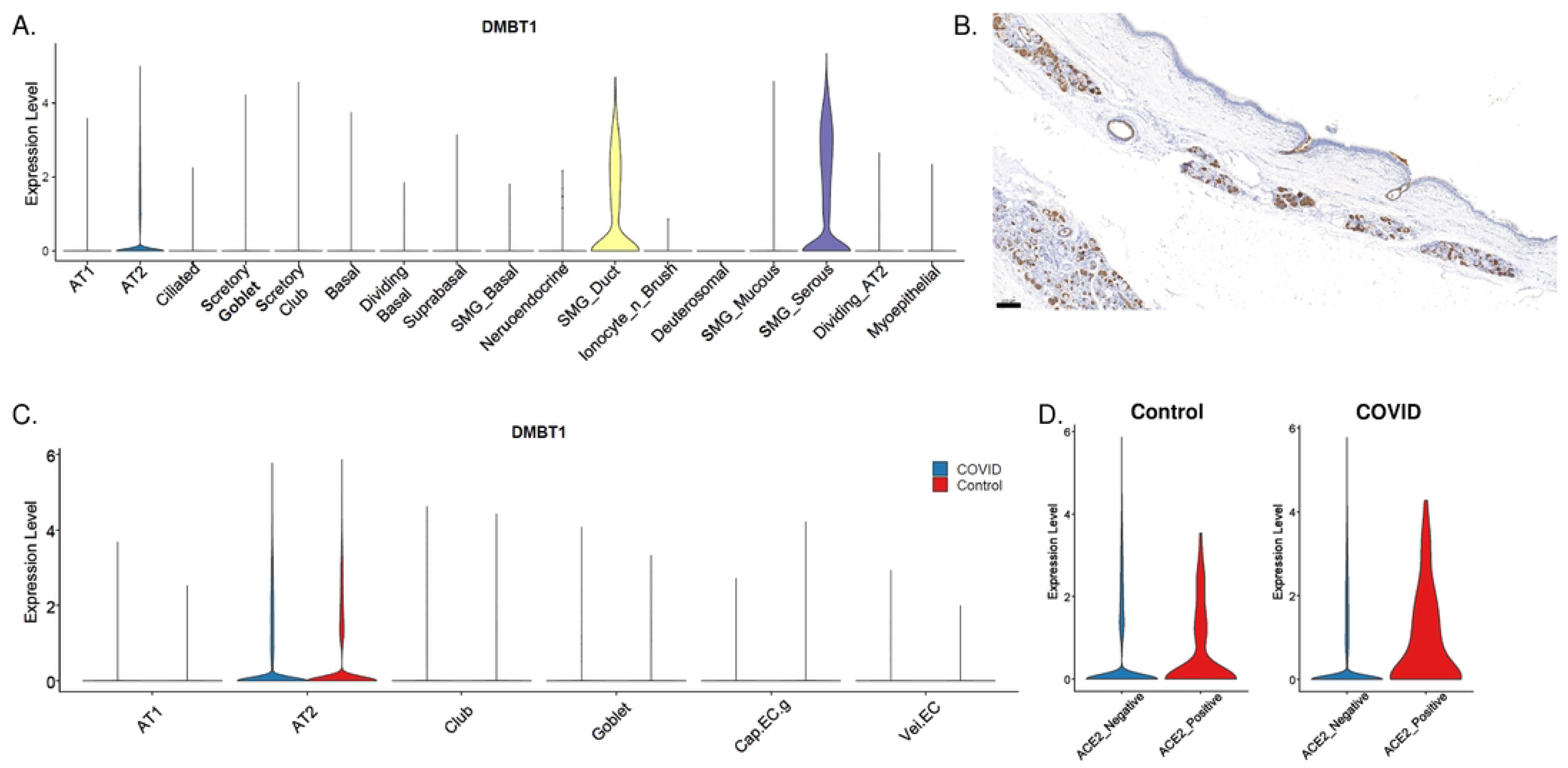
DMBT1 is mainly expressed in the submucosal glands in the respiratory tract. (A) RNA expression of DMBT1 in airway epithelial cell on violin plot from scRNA-seq/snRNA-seq. (B) Section of human trachea stained with the anti-DMBT1 Rabbit Polyclonal antibody. Scale bar = 200 μm. DMBT1 expression was upregulated in ACE2+ AT2 cells of COVID group. (C) RNA expression of DMBT1 in airway epithelial cell of COVID (left blue bars) and control (right red bars) group on violin plot. (D) RNA expression of DMBT1 in ACE2-AT2 cells (left blue bars) and ACE2+ cells (right red bars) on violin plot.

Immunohistochemical staining was performed on human tracheal paraffin-embedded sections using an anti-DMBT1 rabbit polyclonal antibody (Fig 2B), confirming that DMBT1 is predominantly expressed in the respiratory submucosal glands, which may serve as the targets for coronavirus infection.

We further analyzed the correlation between the expression of DMBT1 and the ACE2 receptor in COVID-19[38]. The expression of DMBT1 in AT1 cells, Club cells, Goblet cells, and vein endothelial cells was upregulated in the COVID group compared to the control group, and downregulated in AT2 cells and general capillary endothelial cells (cap.EC.g) (Fig 2C). We categorized the subpopulations of cells into ACE2-Negative (ACE2^-^) and ACE2-Positive (ACE2^+^) groups. Notably, in the COVID group, the expression of DMBT1 in ACE2^+^ AT2 cells was higher than that in ACE2^-^ AT2 cells (Log2FC = 0.614), whereas no significant difference in DMBT1 expression was observed between ACE2^+^ and ACE2^-^ AT2 cells in the control group (Fig 2D). Co-expression of DMBT1 and ACE2 support the role of DMBT1 in SARS-CoV2 infection *in vivo*.

### Peptides of the SRCR domain of DMBT1 can inhibit SARS-CoV-2 infection

Based on the number of conserved cysteine residues in the SRCR domain, the SRCR protein was classified as group A and group B, with 6 cysteines in group A and 8 cysteines in group B. DMBT1 protein belongs to group B SRCR protein and is considered to play roles in tumor suppression and host defense by binding to pathogens. In previous studies[25], the minimal bacterial binding site of the peptide SRCRP2 (QGRVEVLYRGSWGTVC), was identified by using 16 overlapping peptides. We synthesized these 16 overlapping peptides based on SRCR domain (Fig 3A), and used them to block the infection of 293T-DMBT1 cells by authentic SARS-CoV-2 (Fig 3B). The virus was pre-incubated with or without the peptides before adding to 293T-DMBT1 cells. The results showed that peptide 4,5,6,7,9,10 could efficiently inhibit virus infection. Peptides 7 and 9 showed most potent inhibitory effects, with infection rates reducing from around 50% to 15.0%. At lower MOI of 0.6, a similar trend was observed (Fig 3C), with peptide 4, 5, 6, 7, 9, and 10 also contributed to a reduction in viral infection. However, despite the inhibitory effects of peptides 7 and 9 on viral infection, peptide 8, referred to previously as SRCRP2 in bacterial Spy0843 protein inhibition[26], did not exhibit any inhibitory effect.

**Fig 3.**
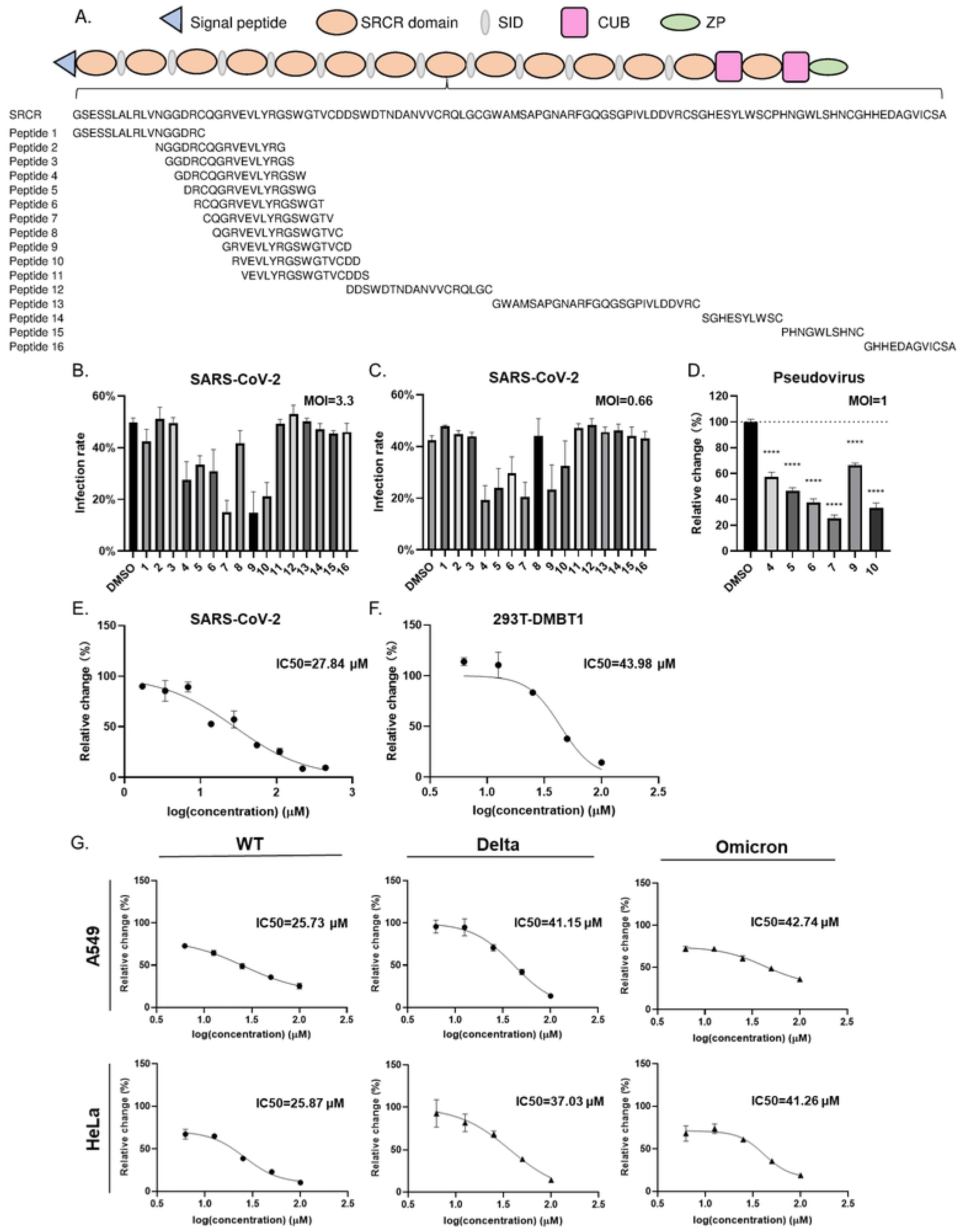
Inhibition of SARS-CoV-2 infection by SRCR peptides derived from DMBT1. (A) Schematic diagram of DMBT1 protein structure. Full length amino acid sequence of SRCR domain of DMBT1 and overlapping peptide library of SRCR domain. (B) Infection rate at SARS-CoV-2 virus with MOI at 3. (C) Infection rate at SARS-CoV-2 virus with MOI at 0.6. (D) Inhibition of peptide of SARS-CoV-2 pseudovirus infection. (E-G) IC50 values were determined using logistic regression analysis. Relative change rate was calculated based on comparison to DMSO control. (E) 293T-DMBT1 was infected by SARS-CoV-2 original strain (SH01) after peptide treatment. Virus incubated with peptide 7 in a 1:2 gradient dilution from 800 μg/mL, and then added to cells. Detection of viral infection by immunofluorescence at 48 hpi. (F) 293T-DMBT1 was infected by SARS-CoV-2 Delta pseudovirus after peptide treatment. (G) HeLa and A549 cells were infected by SARS-CoV-2 pseudovirus 3 strains after peptide treatment. (F, G) Pseudovirus (MOI=5) was incubated with peptide 7 in a 1:2 gradient dilution from 100 μM, and then added to cells. Intracellular Fluc activity of cell lysates was determined at 48 hpi.

We further tested the inhibitory effect of DMBT1 peptides in 293T-DMBT1 cells infected by pseudovirus bearing the spike protein of original SARS-CoV-2. At a peptide concentration of 0.2 mg/mL, the pseudovirus infection treated with peptide 4, 5, 6,7, 9 and 10 was significantly reduced compared to the DMSO control group (Fig 3D). The addition of peptide 7 reduced viral infection by 75%. Furthermore, we serially diluted the peptides to determine the IC50 values. Peptide 7 exhibited the strongest antiviral effect, and the IC50 of peptide 7 in inhibiting the infection of 293T-DMBT1 cells by authentic SARS-CoV-2 was determined as 27.84 μM (Fig 3E). The IC50 in blocking Delta strain pseudovirus infection was also determined as 43.98 μM (Fig 3F).

We further determined whether the peptides of the SRCR domain inhibit the SARS-CoV-2 pseudovirus infection in wild-type cells including HeLa and A549 cells (Fig 3G and Supplementary Fig 1). Peptides 7 and 9 were co-incubated with SARS-CoV-2 pseudoviruses (MOI = 5; WT, Delta, Omicron strains) at 37°C for 1 h before cell infection. Luminescence (RLU) was measured after 48 hours. Peptide 7 showed the strongest antiviral effect (Fig 3G). At the concentration of 100 μM, viral infection was reduced by around 80%. Peptide 7 inhibited the infection of A549 cells by the WT, Delta, and Omicron strains of SARS-CoV-2 pseudovirus, with IC50 values of 25.73 μM, 41.15 μM, and 42.74 μM, respectively. It also inhibited the infection of HeLa cells by these three pseudoviruses with IC50 values of 25.87 μM, 37.03 μM, and 41.26 μM, respectively.

### Peptide 7 inhibits SARS-CoV-2 binding and internalization but has limited effect on viral release

To investigate the antiviral mechanism of peptide 7, we tested its inhibition in virus binding, internalization, and release steps. We found that preincubation of the virus with peptide 7 reduced the attachment of SARS-CoV-2 to A549-ACE2 cells at 4°C (Fig 4A). The binding of the virus preincubated with peptide 7 was reduced by 50% when compared to peptide 16 that lacks antiviral efficacy. Following binding, the cultures were shifted to 37°C for 1 h to allow virus internalization, after which the viral RNA in cell lysates from peptide 7-pretreated group was significantly decreased (Fig 4B), amounting to only 35% of the control peptide 16. However, pretreatment of A549-ACE2 cells with peptide 7 for 1 hour before SARS-CoV-2 infection did not significantly inhibit the binding and internalization (Supplementary Fig 2). This suggests that peptide 7 primarily targets the SARS-CoV-2 virus rather than the host cells.

**Fig 4.**
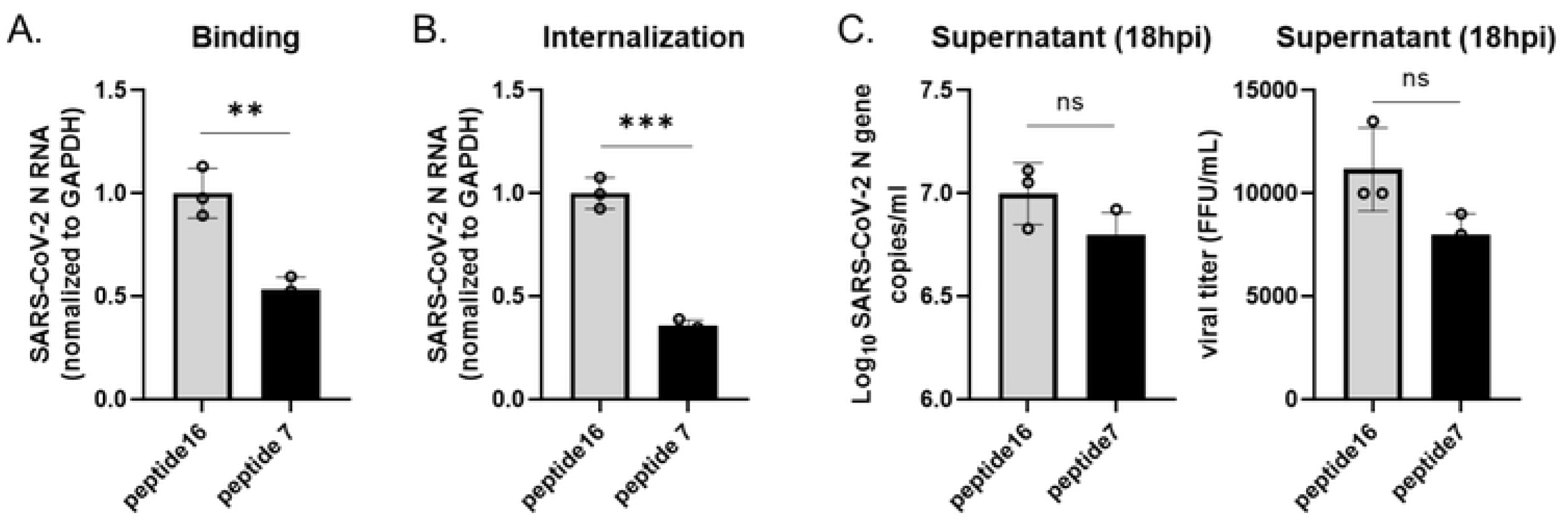
Peptide 7 inhibits SARS-CoV-2 binding and internalization but has limited effect on viral release. (A) Peptide 7 significantly inhibits the binding of SARS-CoV-2 to A549-ACE2 cells. Peptide 7 (100 μg/mL) was pre-incubated with the virus for 1 hour before infecting A549-ACE2 cells (MOI=10). Viral RNA in cell lysates was detected by qPCR, normalized to GAPDH. P values were calculated by comparison with the peptide 16. (B) Peptide 7 significantly inhibits the internalization of SARS-CoV-2 into A549-ACE2 cells. (C) Peptide 7 has limited effect on SARS-CoV-2 release from A549-ACE2 cells. A549-ACE2 cells were infected with SARS-CoV-2 (MOI=0.01) for 14 hours and then treated with peptide 7 or 16 (150 μg/mL) for 18 hours. The supernatant was then collected at 18 hpi for qPCR and FFA analysis. P values were calculated by comparison with the peptide 16.

To evaluate the effect of peptide 7 on viral release, A549-ACE2 cells were infected with SARS-CoV-2, then peptide 7 or peptide 16 was added to culture medium at 14 hours post-infection (hpi), and the supernatant was collected 18 hpi. qPCR and focus forming assay (FFA) showed that peptide 7 did not significantly inhibit viral release (Fig 4C).

### Peptide7 can bind to Spike trimer of SARS-CoV-2

To study the mechanism by which peptide 7 inhibits SARS-CoV-2 infection, we tested its direct binding to virus spike trimer protein. We also tested its binding to host viral receptor ACE2. Binding of spike or ACE2 protein to plate-bound peptide 7 was measured by ELISA. The binding results showed that peptide 7 can bind to both the spike and the ACE2 protein (Fig 5A and 5B), whereas the control peptide 16 lacks binding capacity. Furthermore, peptide 7 does not interfere with the binding between Spike and ACE2 (Fig 5C).

**Fig 5.**
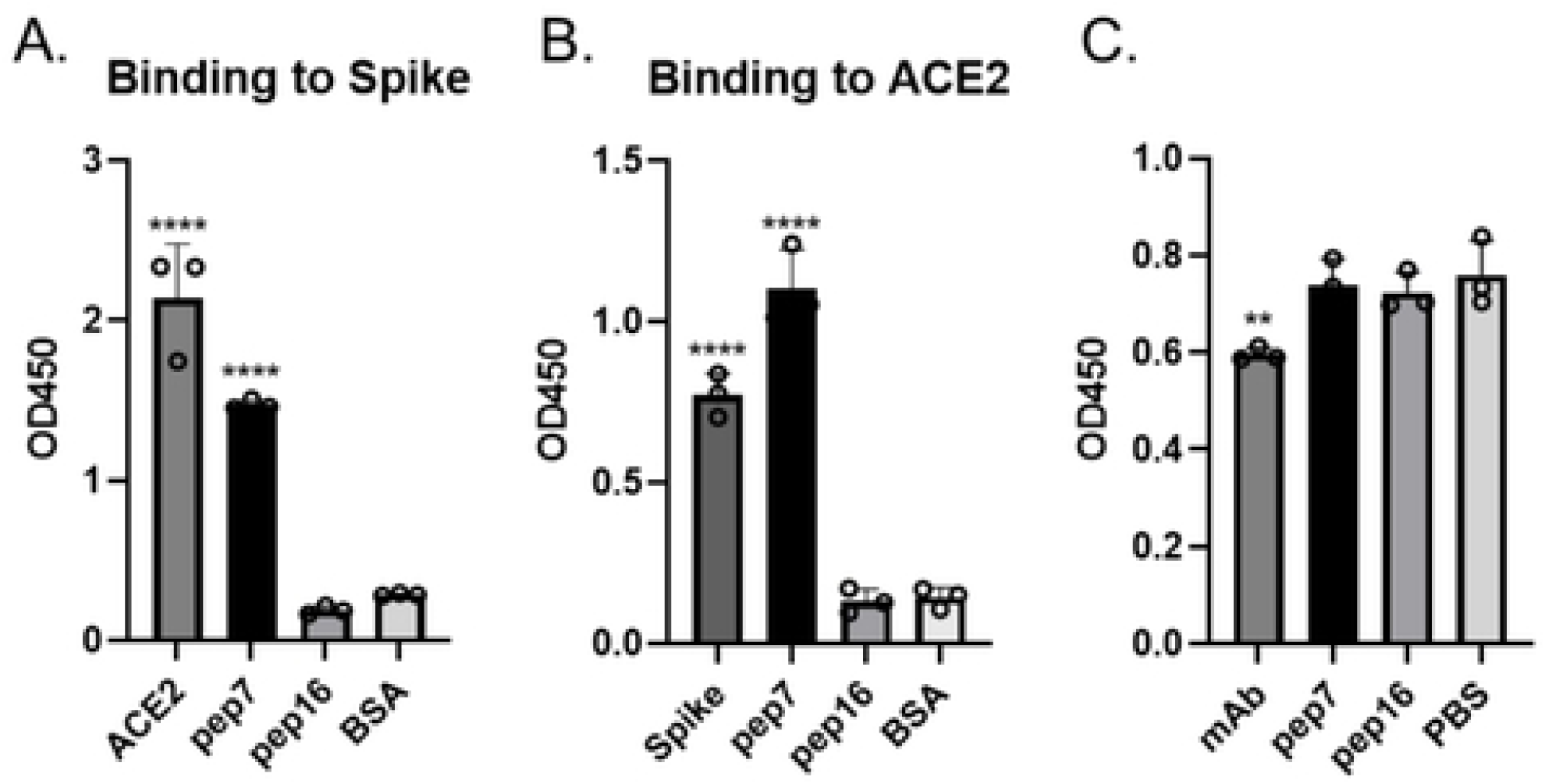
Peptide 7 can bind to Spike trimers and ACE2. (A) Binding of spike trimers with plate coated peptides (1 μg/well), BSA (negative control), or ACE2 protein (positive control, 100 ng/well). (B) Binding of ACE2 with plate coated peptides (1 μg/well), BSA (negative control), or spike protein (positive control, 100 ng/well). ****P<0.0001 when compared with BSA. (C) Influence of Pep7 on the binding of ACE2 to Spike trimer: binding of ACE2 to plate-coated spike was measured by anti-ACE2 antibody. Peptides were added to measure the influence on Spike-ACE2 binding. mAb (Spike Neutralizing Antibody, Clone 40591-MM43) was used as a positive control in blocking Spike-ACE2 binding. **P < 0.01 when compared with PBS.

### Peptide 7 inhibits SARS-CoV-2 infection in hamster model

We then tested the *in vivo* efficacy of peptide 7 in inhibiting SARS-CoV2 infection using the Syrian hamster model. Hamsters were infected with SARS-CoV-2 wild-type strain (Wuhan-01) after co-incubation with peptides. Animals inoculated intranasally (i.n.) with untreated virus showed significant weight loss as early as 2 days post-infection, with about 25% weight loss by 7 days post-infection (Fig 6A). Compared to the untreated control group, the high-dose peptide 7 treatment group protected animals from weight loss. The low-dose peptide 7 treatment group also reduced the extent of weight loss, with animals regaining weight by 4 days post-infection, earlier than the untreated control group, and a significant difference was observed by 5 days post-infection. RT-qPCR analysis of viral genomic RNA in the lungs at 4 days post-infection revealed a 68% reduction in RNA expression (*P* < 0.005) in the high-dose peptide 7 group compared to control group of peptide 16, indicating a significant decrease in SARS-CoV-2 particles in the lungs (Fig 6B). We analyzed lung specimens from the necropsy group at 4 dpi and calculated the pathology score (Fig 6C). All lungs from the untreated and irrelevant peptide control groups exhibited similar histological symptoms of pneumonia, including extensive alveolar septal thickening, inflammatory cell infiltration, hemorrhage, exudate in vessels or alveolar spaces and a small amount of syncytium (Supplementary Fig 3). The animals treated with high-dose peptide 7 group had significantly lower lung histological indications (Fig 6D). The viral load in the lungs of animals treated with high-dose peptide 7 group was significantly reduced, along with prevention of weight loss and lung pathological changes. The high-dose peptide 7 treatment significantly mitigated pulmonary inflammation and lesions, with pathological manifestations characterized by milder inflammation, multifocal consolidation, slight thickening of the alveolar walls, minimal monocyte infiltration, and the absence of alveolar hemorrhage.

**Fig 6.**
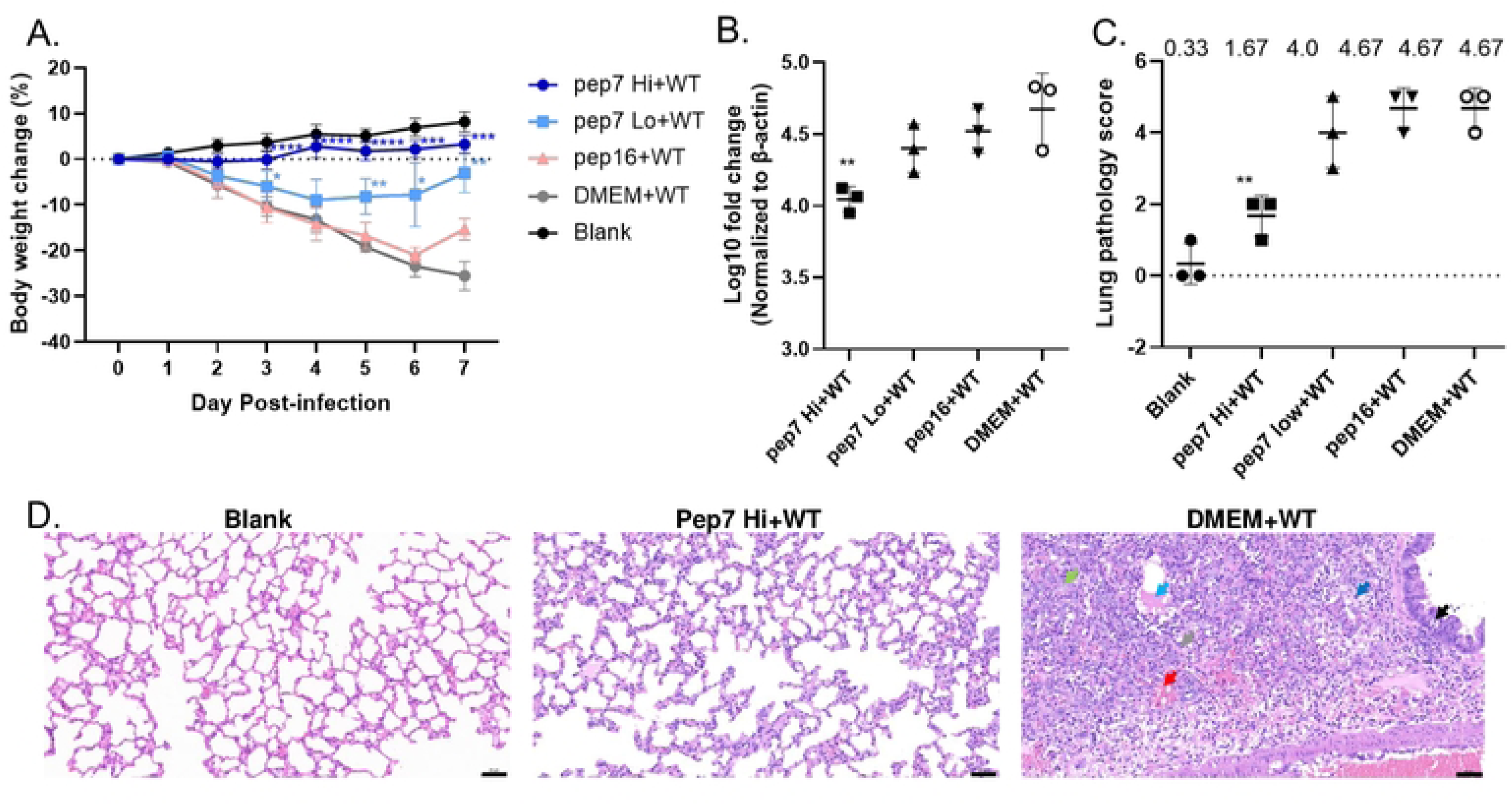
Peptide 7 inhibits SARS-CoV-2 infection in hamsters. (A) The body weight of WT strain-infected hamsters treated with peptide 7(high dose: 600 μg/hamster, low dose: 100 μg/hamster), peptide 16(600 μg/hamster), DMEM, or blank. Hamster body weights were recorded daily at 0-7 dpi (n=3 per group). Weight loss was defined as percentage loss from 0 dpi. Statistical significance of daily body weight comparisons between peptide-treatment groups and the DMEM + WT group was analyzed using unpaired t-tests with GraphPad Prism 8. (B) Analysis of the virus load in the lung at 4 dpi (n=3 per group). Data represent log10fold change normalized to β-actin (relative to Blank group). Statistical significance was determined by one-way ANOVA using DMEM+WT group as the reference using GraphPad Prism 8. (C) Lung pathology scoring in each group. Statistic analysis was performed with the DMEM+WT group as control by one-way ANOVA using GraphPad Prism 8. (D) Representative histopathology of the lungs from blank group, peptide 7 high dose group, or DMEM group hamsters at 4 dpi. Granulocytes (blue arrow), necrotic cell fragments (gray arrow), eosinophilic tissue fluid (light blue arrow), alveolar hemorrhage (red arrow), thickening of alveolar septa (light green arrow). Scale bar = 50 μm. Data were representative of three independent experiments.

### The cysteine of peptide 7 is a key residue for dimerization and antiviral activity

During the course of our research, we observed that peptide 7 from earlier synthetic batches spontaneously formed homodimers in PBS. Despite this dimerization, the antiviral efficacy of peptide 7 was not significantly reduced (Supplementary Fig 5). We hypothesized that this spontaneous homodimer formation might be due to the formation of intermolecular disulfide bonds between the free cysteine residues at the N-terminus. To test this hypothesis, we chemically synthesized a disulfide-linked homodimer of peptide 7 (Pep7-dimer, purity >90%) and evaluated its antiviral activity. The results demonstrated that the IC50 values (based on mass concentration) of Pep7-dimer for inhibiting both live virus and pseudovirus infections showed no significant difference compared to monomeric peptide 7 (Fig 7A). This finding not only confirmed our hypothesis that disulfide bonds mediate the homodimerization of peptide 7 but also indicated that such dimerization does not significantly affect its antiviral activity.

**Fig 7.**
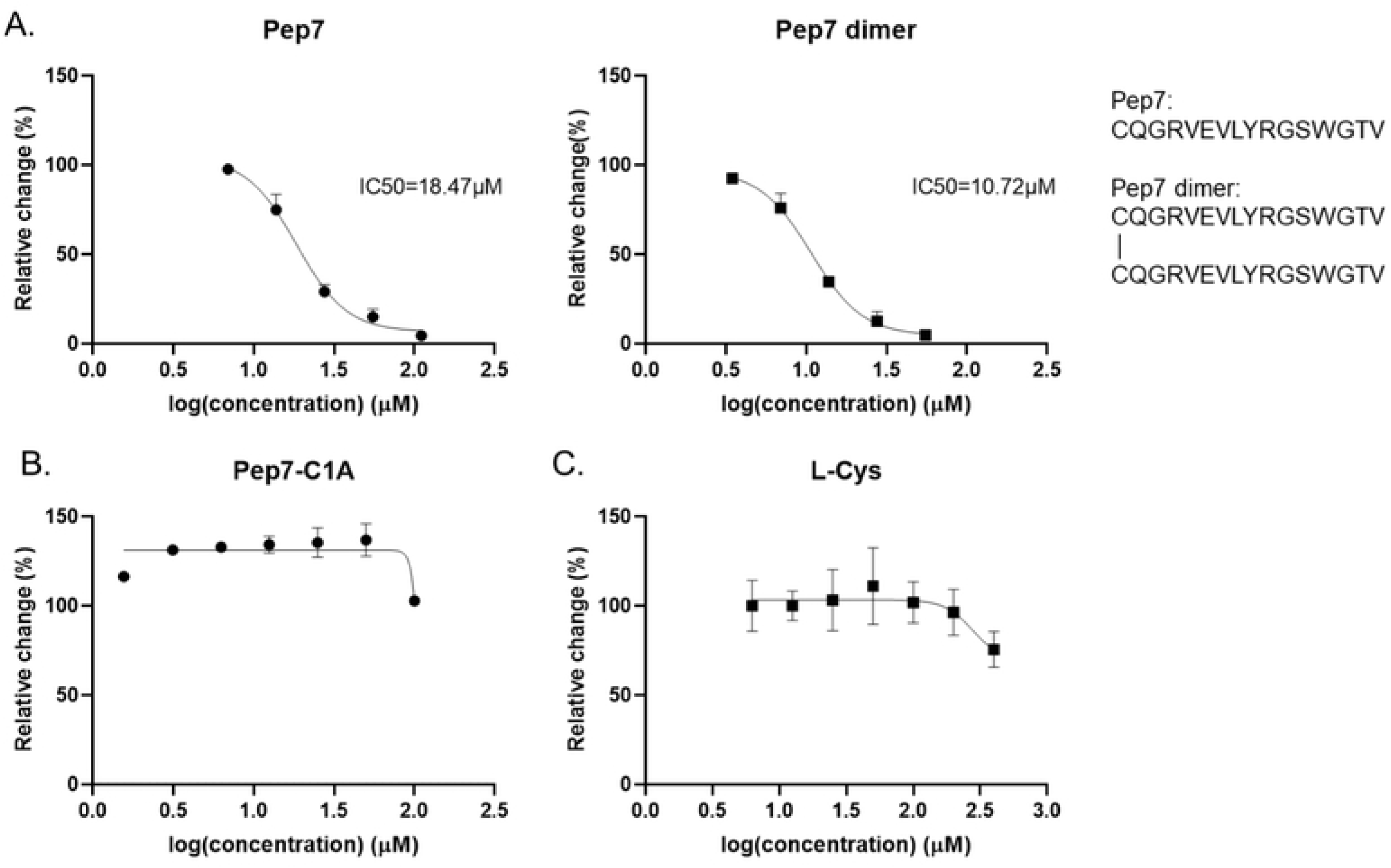
The cysteine of peptide 7 is a key residue for peptide 7 dimerization and antiviral activity. (A) Antiviral activity of peptide 7 and its disulfide-linked homodimer (2Pep7) against pseudovirus infections, as measured by IC50 values based on mass concentration. Pseudovirus incubated with Pep7 or Pep7 dimer in a 1:2 gradient dilution from 200 μg/mL. (B) Replacement with alanine (Pep7 C1A) or (C) free L-cysteine alone completely abolished antiviral activity. Intracellular Fluc activity of cell lysates was determined after 48h of infection. The relative change rate was calculated based on the DMSO control.

To further investigate the critical role of the N-terminal cysteine in the antiviral activity of peptide 7, we synthesized a mutant peptide, Pep7-C1A, by replacing the N-terminal cysteine with alanine, which lacks a thiol group (Fig 7B). Experimental results showed that Pep7-C1A completely lost its antiviral activity. To rule out the possibility that free cysteine itself might exhibit antiviral effects, we conducted experiments using L-cysteine alone and found that even at a high concentration (400 μM), L-cysteine exhibited almost no antiviral activity(Fig 7C). These results indicate that the N-terminal cysteine is not only a key residue for the dimerization of peptide 7 but also an essential residue for its antiviral activity.

## Discussion

### Paradoxical roles of soluble and membrane forms of DMBT1 in SARS-CoV2 infection

As a critical mucosal surface protein, DMBT1 plays a significant role in various viral and bacterial infections. We found that membrane-bound DMBT1 enhances ACE2-dependent SARS-CoV-2 infection. DMBT1 is highly expressed by submucosal glands of human trachea as examined by immunohistochemistry. This is consistent with previous findings that submucosal gland is a target of SARS-CoV-2 infection. Antiviral treatment in submucosal gland might serve as a unique target as scavenger protein for clearance of virus and avoiding long COVID[40, 41].

On the other hand, previous studies have found that DMBT1^GP340^ exhibits antiviral activity against HIV and IAV infection, and this antiviral effect is mediated by the interaction between the virus and the carbohydrates on DMBT1[11-13, 15, 16]. However, in our study, the antiviral efficacy of peptides derived from the SRCR domain of DMBT1 contains no glycosylation sites, highlighting a distinct mechanism of action.

### Molecular mechanisms of SRCR domain in SARS-CoV-2 interaction

The scavenger function of DMBT1 on bacteria was reported to be mediated by SRCR. We have identified peptide 7 (CQGRVEVLYRGSWGTV) within the SRCR domain of DMBT1, which exhibits inhibitory activity against viral infection. This peptide effectively inhibits the infection of authentic SARS-CoV-2 and its pseudovirus, with an IC50 value in the range of 40 micromoles. Additionally, the peptide demonstrates antiviral activity in a Syrian hamster model with reduced viral titers in lung tissues, alleviating weight loss and lung inflammation associated with viral infection.

Furthermore, we identified the N-terminal cysteine residue of peptide 7 as critical for its antiviral activity. Mutation of this residue completely abolished the peptide’s antiviral effects, while free cysteine alone exhibited no activity. This highlights the structural importance of the N-terminal cysteine and provides a foundation for future studies on structural insights, virion-DMBT1 interaction, and drug design. Notably, previous studies have primarily focused on the functional roles of VEVL and W residues, underscoring the novelty of our findings[25, 26, 42]. Molecular simulation of DMBT1 protein containing 14 SRCR domains showed that peptide 7 forms a hairpin-like beta-sheet (Supplementary Fig 4). Binding experiments showed that peptide CQGRVEVLYRGSWGTV can bind to spike protein and ACE2. A similar antiviral mechanism has been reported for peptide 4H30, derived from human β-defensin 2 (HBD2), which also binds to both Spike and ACE2 and inhibits viral internalization and release through aggregation and crosslinking[43].

In conclusion, this study not only reveals the potential of peptide 7 as a SARS-CoV-2 infection inhibitor but also provides new insights into its mechanism of action. These findings lay the groundwork for understanding the viral entry and internalization, as well as the development of DMBT1-based antiviral therapies against SARS-CoV-2 and other emerging viral pathogens. Future research should focus on elucidating the molecular mechanisms underlying the structure basis of SRCR-virion binding, and optimizing inhibitors like peptide 7 to enhance its antiviral efficacy, as a broad-spectrum antiviral agent against enveloped viruses.

## Materials and methods

### Cloning and expression of DMBT1 in 293T cell line

293T and 293T ACE2 knockout (293T-ACE2KO) cell lines were purchased from ABclonal Technology (Wuhan, China). A549 and HeLa cells were from ATCC. All cell lines were cultured in DMEM medium, supplemented with 10% FBS and 1% penicillin/streptomycin. Full length cDNA of DMBT1 gene was purchased from R&D systems, USA (RDC3108). It was cloned into the pLV-GFP-IRES-puro vector. 293T cells were transfected with pLV-DMBT1-GFP-IRES-puro in Lipofectamine 2000 (Invitrogen, USA) according to the manufacturer’s instruction and selected with 2 μg/ml puromycin (Invitrogen, USA). DMBT1-positive cells were sorted by flow cytometry as described below and sorted the dual positive cell population.

### Flow cytometry

DMBT1 expression was detected using 1μg/ml GP340 monoclonal antibody (Invitrogen, USA), and secondary antibody anti-mouse IgG APC (Biolegend, USA) in 1:200 dilution. Cells were detached with 1 mM EDTA (Invitrogen, USA) in PBS without Ca^2+^ or Mg^2+^, and washed three times by centrifugation at 1500 rpm for 3 minutes. Both primary and secondary antibody staining were incubated at 4 °C for 30 minutes.

### Western Blot

Total cellular protein was extracted using RIPA (Beyotime, China) and quantified with the BCA method. For the electrophoresis of DMBT1 protein, a 6% SDS-PAGE resolving gel along with a 250 kDa Protein Ladder (Invitrogen, USA) was employed, whereas for ACE2 and GAPDH proteins, an 8% SDS-PAGE resolving gel combined with a 180 kDa Protein Ladder (Invitrogen, USA) was utilized. Equal volumes of 10 μg samples were subjected to electrophoresis and traditional wet transfer onto PVDF membranes (Millipore, USA) in a Mini-PROTEAN Tetra electrophoresis system (Bio-Rad, USA). The PVDF membranes were then blocked with 5% non-fat dry milk in PBST at room temperature for 2 hours. Incubation with Anti-DMBT1 rabbit pAb (Sino Biological, China) and anti-ACE2 rabbit mAb (ABclonal, China) was performed overnight at 4°C. The secondary antibody used was horseradish peroxidase-conjugated goat anti-rabbit IgG (H+L) (Southernbiotech, USA), incubated for 2 hours at room temperature. After each incubation, the membranes were washed three times with PBST. Subsequently, the PVDF membranes were detected using the Immobilon Western HRP substrate (WBKLS0100, Millipore, USA).

### Immunohistochemistry

The DMBT1 protein was detected by using a rabbit polyclonal antibody (Sino Biological, China) at a dilution of 1:1000 and a rabbit polymer detection system (ZSGB-BIO, China). Briefly, paraffin sections were dewaxed with xylene and ethanol gradients to water, and antigen retrieval was performed with citrate antigen retrieval solution (pH 6.0). Tissue sections were incubated with 3% hydrogen peroxide solution at room temperature in the dark for 25 min to abolish endogenous peroxidase activity, and blocked in 3% BSA in PBS for 30 minutes at room temperature. The sections were then incubated with the primary monoclonal antibody overnight at 4°C. The HRP labeled Goat anti rabbit IgG antibody was then applied for 20 minutes at 37°C. The reaction product was visualized by using 3-3′-diaminobenzidine tetrahydrochloride, counterstained with Harris hematoxylin, and sealed with neutral resin after dehydration.

### Infection by transcription and replication-competent SARS-CoV-2 virus-like-particles

Transcription and replication-competent SARS-CoV-2 virus-like-particles (trVLP-Nluc, based on nCoV-SH01, GenBank accession no. MT121215) was generated using a protocol as we previously described[32, 33]. The single-cycle trVLP-Nluc particles, in which the N gene is replaced by NanoLuc luciferase reporter, were packaged in N-trans-complemented cells. 96 well plates were coated with Poly-D-lysine before host cells were seeded. Host cells were infected with trVLP-Nluc at indicated dose and incubated at 37 °C with 5% CO_2_ for 24 hours. The nano-luciferase activity was determined using Nano-Glo® Luciferase Assay kit (N1110, Promega, USA) according to the manufacturer’s instructions.

### Infection by pseudovirus

Host cells were seeded in 96 well plates (30000 cells per well). HIV-1-based pseudoviruses (Novoprotein, China) encoding SARS-CoV-2 spike variants (D614G, Delta, and Omicron) and a luciferase reporter gene were generated in 293T cells. Pseudovirus was added in final volume of 150 μL. Infected host cells were cultured at 37 °C with 5% CO_2_ for 48 hours. The luciferase activity was determined using ONE-Glo Luciferase Assay kit (E6110, Promega, USA) according to the manufacturer’s instructions.

### Blocking of virus infection by overlapping peptides of SRCR domain of DMBT1

Peptides of SRCR domain of DMBT1 were synthesized by Wuxi AppTec, China. SARS-CoV-2 pseudovirus was co-incubated with serially diluted peptides in culture medium at 37 °C for 1 hour. After incubation, 100 μL peptide-pseudovirus mixture was added to host cells, and the final volume was 150 μL. Cells were incubated at 37 °C with 5% CO_2_ for 48 hours. The measurement of infection by luciferase activity was described above.

To measure the blocking efficacy of peptides for SARS-CoV-2 authentic virus, peptides were incubated with SARS-CoV-2 original strain (SH01) (MOI 3 and 0.6) for 30 minutes at room temperature[34]. The concentration of each peptide was 0.2 mg/ml. Host cells were inoculated with virus mixed with blocking peptides. After 24 hours, cells were fixed with 4% PFA in PBS for immunofluorescence staining and high-content analysis as described[35]. In brief, SARS-CoV-2 virus-infected host cells were washed twice with PBS, fixed with 4% PFA in PBS for 30 minutes, then permeabilized with 0.2% Triton X-100 for 1 hour. Cells were then incubated with house-made mouse anti-SARS-CoV-2 nucleocapsid protein serum (1:1000) at 4 °C overnight. After three washes, cells were incubated with the goat anti-mouse IgG (H+L) secondary antibody conjugated with Alexa Fluor 555 (2 μg/ml, A21424, Invitrogen, USA) for 2 hours at room temperature, followed by staining with 4′,6-diamidino-2-phenylindole. Images were collected using an Operetta High Content Imaging System (PerkinElmer), and processed using the PerkinElmer Harmony high-content analysis software v4.9 and ImageJ v2.0.0.

### *In vivo* antiviral efficacy of peptide 7 in hamster model

The SARS-CoV-2 original strain (Wuhan-01) was isolated from a laboratory-confirmed COVID-19 patient in Vero E6 cells[36]. All animal experiments involving infectious virus were conducted in a biosafety level 3 (BSL-3) facility at the Second Military Medical University. Female hamsters (6-8 weeks old) were housed in a BSL-3 laboratory with ad libitum access to standard pellet feed and water.

To evaluate the antiviral activity of the peptides, peptide 7 was mixed with SARS-CoV-2 virus (100 μL in culture medium, 6×10^5^ PFU/hamster) at high-dose (600 μg per hamster), or low-dose (90 μg per hamster). Negative control peptide 16 (600 μg per hamster), DMSO, or DMEM containing 2% FBS were incubated separately with SARS-CoV-2 virus (100 μL, 6×10^5^ PFU/hamster) at 37°C for 1 hour, then intranasally inoculated into hamsters. The body weight of the hamsters was recorded daily. Lung tissue was collected at 4 days post-infection, and tissue pathology was measured by H&E staining. Total RNA was extracted from lung homogenates to determine viral titers using qPCR. In brief, total RNA was extracted from lung tissue using TRIzol reagent (Invitrogen, USA). RNA was then converted into cDNA using a reverse transcription system (Vazyme, China). The cDNA products were directly used for subsequent qPCR analysis with Taq Pro Universal SYBR qPCR Master Mix (Vazyme, China) and gene-specific primers. Hamster β-actin expression was utilized for normalization. Primers and probes used are as follows: CoV-N-Fwd: 5′-AAGGCGTTCCAATTAACACCA-3′; CoV-N Rev: 5′-TGCCGTCTTTGTTAGCACCA-3′; hamster β-actin-Fwd: 5′-AGCAGTCTGTTGGAGCAAGC-3′; hamster β-actin-Rev: 5′-TCTAGGGAATTGGGGTGGCT-3′.The SARS-CoV-2 viral load was quantified by RT-qPCR and analyzed using the 2^-ΔΔCt^ method[37]. Data are presented as mean ± standard deviation (SD) and were analyzed using unpaired t-tests with GraphPad Prism 8.0. A p-value of less than 0.05 was considered statistically significant.

### Binding and internalization of SARS-CoV-2

For the binding and internalization assays, plates were coated with poly-D-lysine, and cells were seeded to achieve 100% confluence. For the virus binding assay, the SARS-CoV-2 virus (SH01) was diluted in DMEM containing 2% FBS to an MOI of 10 and pre-incubated with peptide 7 or peptide 16 (100 μg/mL) at 37°C for 1 hour. Both the virus-peptide mixture and cells were pre-chilled on ice, after which the medium was replaced with the ice-cold virus mixture and incubated on ice for 45 minutes. Unbound virions were removed by washing the cells 5 times with ice-cold DMEM containing 2% FBS. Cells were lysed to extract total RNA with the TRIzol reagent (Invitrogen, USA), and bound viral RNA was quantified by RT-qPCR using the One Step PrimeScript™ RT-PCR Kit (RR064B, TaKaRa) on CFX Connect Real-Time System (Bio-Rad, California, USA).

For the virus internalization assay, after binding and washing, cells were incubated with 2% FBS DMEM at 37°C for 1 hour to allow virus internalization, followed by incubation on ice for at least 5 minutes. Cells were treated with ice-cold proteinase K (400 μg/mL in PBS) for 45 minutes to remove surface-bound virions, followed by washing with ice-cold 2% FBS DMEM. Cells were lysed to extract total RNA, and internalized viral RNA was quantified by RT-qPCR. The relative RNA abundance of bound or internalized virions was normalized to internal control GAPDH. Primers and probes used are as follows: nCoV-N-Fwd: 5′-GACCCCAAAATCAGCGAAAT-3′; nCoV-N Rev: 5′-TCTGGTTACTGCCAGTTGAATCTG-3′; nCoV-N-Probe: 5′-FAM-ACC CCGCATTACGTTTGGTGGACC-BHQ1-3′; hGAPDH-Fwd: 5′-TGCCTTCTTGCCTCTTGTCT-3′; hGAPDH-Rev: 5′-GGCTCACCATGTAGCACTCA-3′; and GAPDH-Probe: 5′-FAM-TTTGGTCGTATTGGGCGCCTGG-BHQ1-3′.

### Virus titer determination by focus-forming assay

The experiment was per formed similarly as described previously[34]. Briefly, Vero E6 monolayer in 96-well plates was inoculated with serially diluted virus for 2 h and then overlaid with methylcellulose for 48 h. Cells were fixed with 4% PFA in PBS for 1h and permeablized with 0.2% Triton X-100 for 1h. Cells were stained with home-made mouse serum against N protein of SARS-CoV-2 overnight at 4°C, incubated with the secondary antibody HRP-conjugated goat anti-rabbit IgG (H+L) (Invitrogen, USA) for 2 h at room temperature. The focus-forming unit was developed using TrueBlue substrate (5510-0030, Sera Care, USA).

### Hematoxylin and Eosin (H&E) Staining

Hamster lung tissues were preserved in 4% paraformaldehyde and embedded in paraffin. Tissue sections (3 μm) were dewaxed and rehydrated, followed by conventional H&E staining. The full images of tissue sections were taken by Pannoramic MIDI (3DHISTECH, Hungary). Based on H&E staining scans, syncytia, alveolar septal thickening, inflammatory cell infiltration, alveolar exudate, and hemorrhage were annotated.

### Peptide-binding assay

Peptide binding to Spike trimers were measured with recombinant ACE2 as positive control. Peptides (1 μg/well), ACE2 (100 ng/well, Sinobiological, China), or BSA as negative control, were dissolved in PBS and coated onto the ELISA plate and incubated overnight at 4°C, followed by overnight blocking with 2% BSA at 4°C. Then the Spike trimer was added to the coated wells and incubated at 37°C for 1 hour. The binding to Spike trimer was detected by incubating with mouse anti-Spike antibody (40591-MM43, 1 μg/ml, Sinobiological, China) for 50 minutes. The reaction was developed by adding 100 μL of TMB solution (Biolegend, USA) at 37°C for 15 minutes and terminated with 50 μL of 1M H_2_SO_4_. The readings were obtained using a microplate reader (Spark, Tecan, Switzerland) at 450 nm. Data are presented as mean ±standard deviation (SD) and were analyzed using One-way ANOVA with GraphPad Prism 8.0.1.

Peptide binding to ACE2 were measured with recombinant Spike trimer as positive control. Peptides (1 μg/well), Spike trimer (100 ng/well, Novoprotein, China), or BSA as negative control, were dissolved in PBS were coated onto the ELISA plate and incubated overnight at 4°C, followed by overnight blocking with 2% BSA at 4°C. Then ACE2 was added to the coated wells and incubated at 37°C for 1 hour. The binding was detected by incubating with rabbit anti-ACE2 mAb (1:2000, ABclonal, China) for 50 minutes. The reaction was visualized and quantified as described above.

### Peptide inhibition of ACE2 binding to Spike trimer

To determine the efficacy of peptides in inhibiting the binding of ACE2 to the Spike protein, peptides or PBST were added to plate-coated Spike trimer and incubated at 37°C for 1 hour, followed by the addition of ACE2 to the wells and further incubation at 37°C for 1 hour. The bound ACE2 was assessed by incubating with rabbit anti-ACE2 antibody for 50 minutes at 37°C, followed by incubation with goat anti-rabbit IgG(H+L)-HRP (1:5000, ABclonal, China) at 37°C for 50 minutes. The reaction was visualized and quantified as described above.

### Sources of single-cell sequencing data

The processed single-cell sequencing data used in this study was derived from the original data from previous studies[38, 39]. The accession numbers for raw snRNA-seq and bulk RNA-seq data from the lungs of COVID-19 patients and controls have been uploaded and deposited by the authors in the Genome Sequence Archive for Human at the National Genomics Data Center (https://ngdc.cncb.ac.cn/gsa-human/), with the respective accession numbers HRA000615, HRA000646, and HRA001136^33^. Before initiating this study, we obtained consent from the authors of the original article to access and utilize these data for subsequent analysis. The scRNA-seq and snRNA-seq data of the tissue architecture of lung and airways can be accessed and downloaded at https://www.lungcellatlas.org/[39].

### Data Processing and Analysis

The raw matrix was loaded using the Seurat R package (version 4.1.1). Batch processing was performed on all samples, and integration was achieved using the RPCA algorithm within Seurat. Gene expression data for the corresponding cells were extracted based on filtered cells in the metadata for cell type annotation. The extracted cell expression matrix was normalized using the LogNormalization method. The FindVariableFeatures function in Seurat was employed to identify a set of highly variable genes, which were then used for principal component analysis (PCA) dimensionality reduction. The first 30 principal components (PCs) determined by the PCElbowPlot function were selected for dimensionality reduction clustering (resolution = 2) and visualization with UMAP or t-SNE. The FeaturePlot and VlnPlot functions were utilized to generate Umap mapping and violin plots, respectively. Differential expression analysis was conducted using the FindMarkers function with a bimodal distribution test (bimod), setting the significance threshold at p-value ≤ 0.05 and |log2FC| ≥ 0.263. Volcano plots and histograms of cell proportions were constructed using the ggplot R package.

## Declarations

### Ethics approval and consent to participate

All animal experiments involving infectious virus were conducted in a biosafety level 3 (BSL-3) facility at Naval Medical University. All experimental protocols adhered to the standard operating procedures approved for BSL-3 animal facilities. No human subjects were involved in this study.

### Consent for publication

All authors have read and agreed to the published version of the manuscript. All the authors are giving consent to publish.

### Availability of data and materials

The authors confirm that the data supporting the findings of this study are available within the article and its supplementary materials.

### Authors’ contributions

C.Z., Conceptualization, Data curation, Formal analysis, Investigation, Methodology, Validation, Writing-original draft, Writing-review and editing.

Z.W., Data curation, Formal analysis, Investigation, Methodology, Validation.

Z.P., Data curation, Formal analysis, Investigation, Methodology.

X.M., Supervision, Methodology.

Y.C., Investigation, Data curation.

W.Z., Software, Visualization.

P.Z., Validation.

H.T., Supervision.

R.Z., Supervision, Validation, Funding acquisition, Writing-review and editing.

D.Z., Conceptualization, Data curation, Formal analysis, Funding acquisition,

Investigation, Methodology, Project administration, Supervision, Validation, Writing-original draft, Writing-review and editing.

### Corresponding authors

Correspondence to Dapeng Zhou; Rong Zhang; Hailin Tang.

## Acknowledgements

We thank Professor Wuyuan Lu from Fudan University for his suggestions on peptide homodimer related experiments.

We are grateful to Wuhan Servicebio Technology Co., Ltd. for providing pathological testing services and to Shanghai Neo Bio-technology Co., Ltd. for providing single-cell sequencing analysis services.

## Supporting information

**Supplementary Fig 1. Peptide 9 block infection of SARS-CoV-2 pseudovirus encoding spike protein from three different viral strains.** HeLa and A549 was infected by SARS-CoV-2 pseudovirus: Wuhan-01 (WT), Delta or Omicron strain co-incubated with peptides. Pseudovirus was incubated with peptides 9 in a 1:2 gradient dilution from 100 μM, and then added to cells. Intracellular Fluc activity of cell lysates was determined after 48h of infection. The relative change rate was calculated based on comparison to DMSO control. These data are supplement to Fig 3.

**Supplementary Fig 2. Peptide 7 pretreatment of cells shows no significant inhibition of SARS-CoV-2 binding and internalization.** Peptide 7 (100 μg/mL) was pre-treated with A549-ACE2 cells for 1 hour prior to SARS-CoV-2 infection (MOI=10). Viral RNA levels in cell lysates were quantified by qPCR and normalized to GAPDH. No significant reduction in viral binding or internalization was observed compared to the control group (n = 3).

**Supplementary Fig 3. Full images of histopathology of hamster lungs (Scale is 1000 μM).** Hamsters were intranasally inoculated with SARS-CoV-2 preincubated with DMEM, peptide 7 or peptide 16. The lung tissues were harvested at 4 dpi for H&E staining and full images were taken by Pannoramic MIDI.

**Supplementary Fig 4. The structure model of the DMBT1 protein and the molecular surface rendered with the electrostatic potential.** (A) The structure model of the DMBT1 protein was predicted using the AlphaFold3 server (https://alphafoldserver.com/), based on the sequence extracted from the NCBI website (protein ID: NP_015568.2). In the cartoon representation, the peptide 7 (CQGRVEVLYRGSWGTV) is highlighted in red, with mutations shown as pink spheres, and disulfide bonds are presented as yellow sticks. (B) The molecular surface rendered with the electrostatic potential. The vacuum electrostatics were generated in PyMOL using default settings. Positive potentials are shown in blue, while negative potentials are shown in red.

**Supplementary Fig 5. Dimerization detection and antiviral effect of peptide 7 in new batch and old batch.** (A) The samples were analyzed using an Agilent HPLC Infinity II 1290 system coupled with a TOF 6230 detector. 20 μL of each diluted sample was injected and separated on a reverse phase column (Agilent AdvanceBio Peptide Plus 2.7 μm 2.1×150mm, PN: 695775-949). Chromatographic separation was achieved using a 5-65% acetonitrile/water gradience containing 0.1% TFA at 40℃, with a flow rate of 0.4 mL/min. (B) A549-ACE2 was infected by SARS-CoV-2 WT pseudovirus after new batch and old batch peptides treatment. Intracellular Fluc activity of cell lysates was determined at 48 hpi.

**Supplemental Movie 1. Structural modeling of DMBT1.** (A) The rocking movie was generated using PyMOL 2.5 based on Supplemental Fig 4. All labels and colors remain unchanged. (B) The rocking movie was generated using PyMOL 2.5 based on Supplemental Fig 4. The vacuum electrostatics were also generated in PyMOL with 40% surface transparency.

